# Genetically engineered insects with sex-selection and genetic incompatibility enable population suppression

**DOI:** 10.1101/2021.06.11.448100

**Authors:** Ambuj Upadhyay, Nathan R Feltman, Adam Sychla, Siba R Das, Maciej Maselko, Michael J Smanski

## Abstract

Engineered Genetic Incompatibility (EGI) is a method to create species-like barriers to sexual reproduction. It has applications in pest control that mimic Sterile Insect Technique when only EGI males are released. This can be facilitated by introducing conditional female-lethality to EGI strains to generate a sex-sorting incompatible male system (SSIMS). Here we demonstrate a proof of concept by combining tetracycline-controlled female lethality constructs with a *pyramus*-targeting EGI line in the model insect *Drosophila melanogaster*. We show that both functions (incompatibility and sex-sorting) are robustly maintained in the SSIMS line and that this approach is effective for population suppression in cage experiments. Further we show that SSIMS males remain competitive with wild-type males for reproduction with wild-type females, including at the level of sperm competition.

## Introduction

Arguably the most successful large-scale insect control approach to date is Sterile Insect Technique (SIT) (1, 2). SIT uses irradiation at sub-lethal doses to sterilize laboratory-reared insects prior to their targeted environmental release. Such an approach was successfully applied to eradicate the New World Screwworm (*Cochliomyia hominivorax*) from North America during the mid-to late-1900’s (3, 4). Since then, SIT has been successfully applied for the broad-scale control of tephritid fruit flies (*e.g. Ceratitis capitata, Zeugodacus cucurbitae, Bactrocera tryoni*, and *Anastrepha ludens*), onion maggots (*Delia antiqua*), tse-tse flies (*Glossina* spp.), and several coleoptera and lepidoptera species (1).

Several factors limit the widespread adoption of SIT for more insect pests. Exposing insects to sterilizing doses of irradiation can impact their longevity and mating competitiveness (5) (cite Lance 2000). Sometimes the lethal dose of radiation is sufficiently close to the sterilizing dose of radiation such that achieving 100% sterilization means that the vast majority of the males will not survive (6). Many batch irradiation approaches do not separate males from females prior to release, and this can both decrease the effectiveness (7), and lead to public health concerns when the female insect is a disease vector (8).

Recently, biotechnology-enabled approaches have been developed that have the potential to complement or replace SIT programs, or allow SIT-like programs that target a wider range of species. Integrating an early acting female lethality system to New World Screwworm without impacting male fitness and fecundity has the potential to improve SIT effectiveness by several fold (9). Release of insects with dominant lethality (RIDL), has been successfully demonstrated to suppress target mosquito populations in field trials in the Caribbean islands and South America (10). Precision-guided SIT (pgSIT) utilizes expression of Cas9 nuclease to knock out female survival and male fertility genes prior to release to the environment(11, 12).

Engineered Genetic Incompatibility (EGI) is a strategy for creating bi-directional (male-to-female and female-to-male) mating incompatibility that mirrors the behavior (13–16). EGI strains are homozygous for haplosufficient lethal genes with haploinsufficient resistance genes. Outcrossing to wildtype passes one copy of each to hybrid offspring to drive 100% hybrid lethality. Male and female carrying the same EGI constructs are not sterile but can reproduce with their like-kind with normal fecundity. In proof of concept publications, this is achieved using dCas9-based programmable transcriptional activators (PTAs) that target endogenous genes for lethal overexpression or ectopic expression. Resistance in the EGI strain is conferred by small mutations in the target sites that prevent PTA binding. Importantly, multiple mutually-incompatible EGI lines can be created that are incompatible with wild-type and with each other, providing a way to rationally engineer negatively correlated cross-resistance for the first time (13).

As with other underdominant biocontrol methods, EGI could be used as a threshold-dependent gene drive (15, 17, 18). Alternatively, it could be coupled to a conditional female-lethal construct, such as the tetracycline (Tet)-repressible positive feedback circuit (19–21), to produce a strain that more closely replicates SIT biocontrol agents. Here we demonstrate the successful engineering of such a system in *Drosophila melanogaster*. We quantify the behavior of individual genetic components, perform male mating competition assays, and test the ability of SSIMS flies to suppress a wild-type population in multi-generational laboratory cage trials.

## Results

### Generating and validating SSIMS lines

To generate the SSIMS line, we made a compound stock that contains two female lethal (FL) constructs (22) on the X chromosome and an EGI construct on the 3rd chromosome (Figure 1). The EGI construct comprised a dCas9-VPR programmable transcriptional activator (PTA) targeting the promoter of a critical developmental morphogen gene. We used a *pyramus*-targeting PTA. Throughout the strain-construction process, a mutation in the promoter of the *pyramus* gene was maintained to prevent binding of the PTA to the *pyramus* promoter, and therefore lethal ectopic expression (Supplementary Figure S1). The SSIMS line used the *foxo* promoter to drive the PTA, as this genotype performed especially well in previous studies (13). Following mating scheme outlined in Supplementary Note 1, we were able to generate viable SSIMS flies, which contained both of the visual markers of the FL genotype (GFP expression) and the EGI genotype (red eyes) (Figure 2a).

**Fig. 1.**
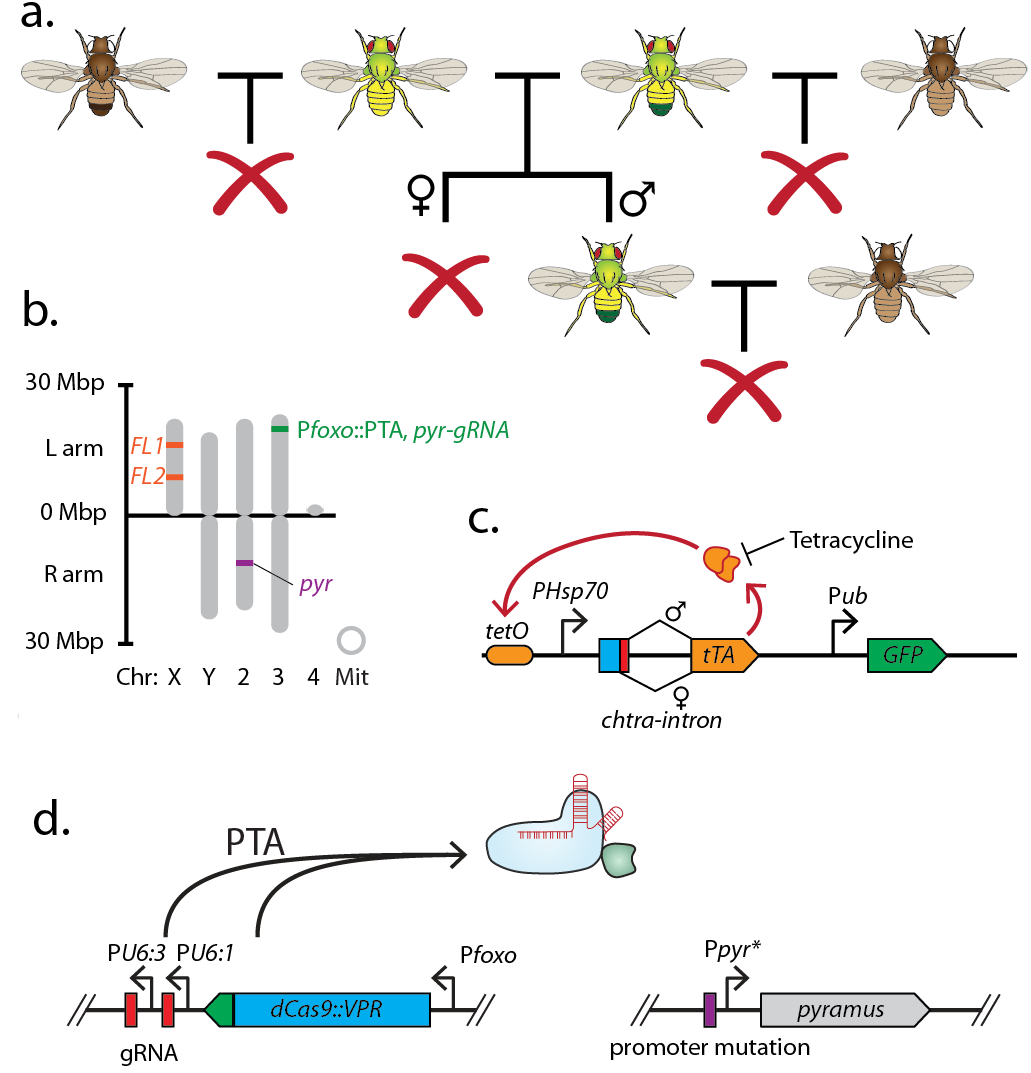
Overall design and implementation of Sex-sorting Incompatible Male System (SSIMS). (a) Illustration of desired behavior of SSIMS insects (green-yellow) when grown in the absence of Tet. Brown flies represent wild-type, and red crosses represent inviable offspring. (b) Genome-scale location of engineered loci in *D. melanogaster* SSIMS line. (c,d) Genetic cassette diagrams for female lethal (FL) and engineered genetic incompatibility (EGI) loci, respectively. SBOL iconography is used for genetic part representation. Abbreviations: FL, X-linked female lethal construct; Mbp, megabasepairs; kb, kilobases; Chr, chromosome; Mit, mitochondrial; PTA, Programmable Transcriptional Activator

**Fig. 2.**
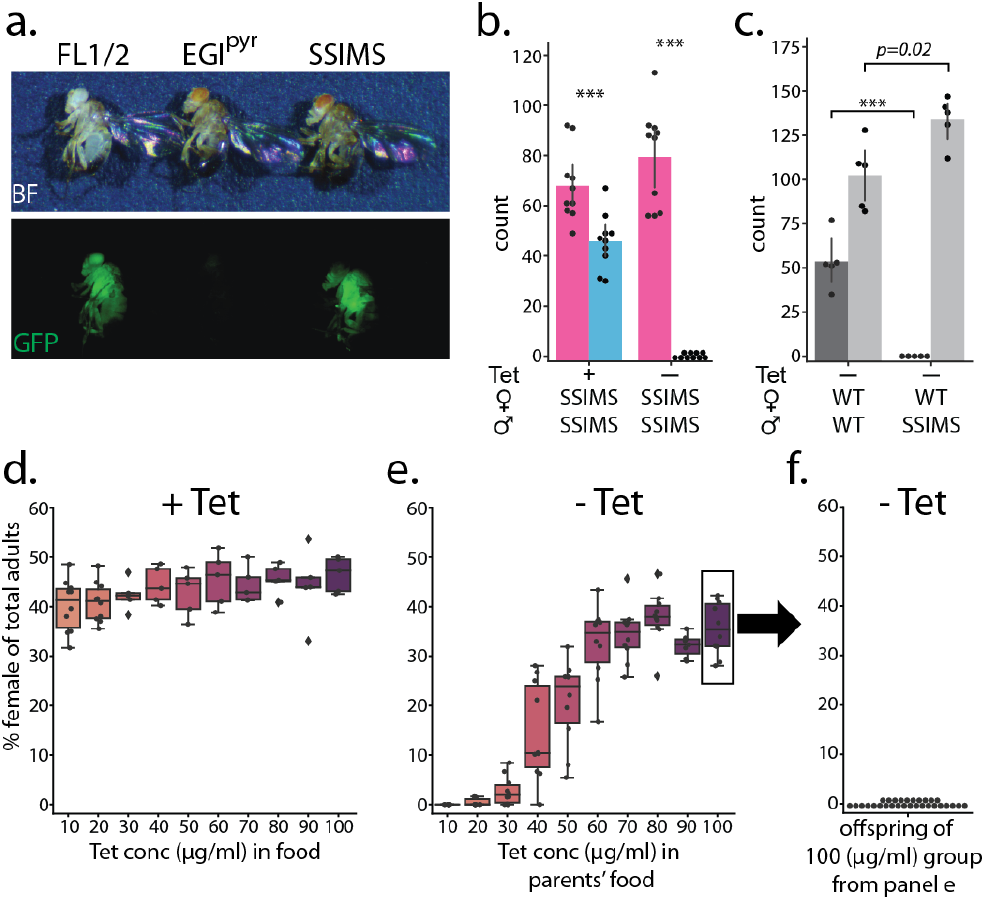
Phenotype of SSIMS flies. (a) Visual markers present in homozygous FL lines (GFP, left) and EGI lines (red eyes, center) are present in SSIMS flies generated via the mating scheme in Supplementary Figure S1. (b) Counts of male (pink) and female (blue) offspring of SSSIMS strain raised in food with 10*μ*g/ml Tetracyline (Tet) and without Tet. No females offspring were observed when larvae are raised in the absence of Tet. Chi-squared test shows significant difference from expected 50:50 ratio of males:female, n=10 per group, ***, *p<0.0001*. (c) Counts of total offspring (dark grey) and egg-lay (light grey) of wild-type females crossed with either wild-type or SSIMS males. SSIMS male mate successfully with wildtype females, producing slightly more eggs, but no adult offspring emerge, n=5 per group, ***, *p<0.0001*. (d) Percent female offspring when SSIMS stocks are reared in food with increasing concentrations of Tet, n = 10 per group. (e) Percent female offspring when respective progeny from (c) are reared in food without Tet, n = 7-10 per group. (f) Percent female offspring when progeny from the 100 *μ*g/ml group in (d) are reared in food without Tet, n = 30. Abbreviations: FL, X-linked Female Lethal; EGI(pyr), engineered genetic incompatibility targeting *pyramus;* WT, wildtype; SSIMS, sex-sorting incompatible male system; Tet, Tetracycline.

### Efficacy of genetic components in SSIMS line

To validate the SSIMS line we first tested whether the two underlying behaviors were maintained in the compound strain (FL and EGI). We confirmed that the SSIMS line bred true in the presence of Tet at 100 *μ*g/ml and contained both visual markers for FL (GFP) and EGI (mini-white) (Figure 2a). In the presence of Tet at (10 *μ*g/ml) the SSIMS line shows a slight sex-ratio bias with fewer females (40% female) surviving to adulthood than expected 50% (Figure 2b). This indicates that the FL transgene is partially leaky in females at 10 *μ*g/ml Tet, and merits further optimization. In the absence of Tet no females survive to adulthood and either die as larvae or pupae, demonstrating the efficacy of the FL transgene (Figure 2b). To confirm the behavior of the EGI components, 20 wild-type females were mated with either 4 wildtype males or 4 SSIMS males, then we quantified number of eggs laid per vial in 24 hours and total number of offspring produced per vial (24 hr egg-lay). Wild-type females did not produce any viable adults when mated with SSIMS male (Figure 2c). We confirmed that SSIMS males successfully mated with wild-type females by counting the number of eggs laid in vials, which showed a slight increase when mated with SSIMS males (Figure 2c).

In an effort to further optimize rearing of SSIMS stock, and minimize leaky FL transgene in females, we titrated Tet in the food and assessed % female in the offsprings both with Tet and when those progeny are transferred to food without Tet (Figure 2d-f). When used in high concentrations, Tet has been reported to carryover between generations by maternal deposition in oocytes (23). SSIMS line grown with 10-100 *μ*g/ml Tet results in 40-45% female (Figure 2d). When these offspring are transferred to food without Tet, their offspring compose of 0-35% female in increasing order correlating with how much Tet the parents were exposed to (Figure 2e). When parents were reared in 100 *μ*g/ml Tet, many of their female offspring survived in the absence of Tet. In these cases, 100% female lethality did not occur until the second generation of rearing on Tet-free media (Figure 2f).

### Male mating competitiveness and evidence for sperm competition

We tested the ability of SSIMS males to compete with wild-type males for mating to wild-type females using a fecundity assay (Figure 3a,c). Twenty virgin females were mated with four males at a 4:0, 3:1, 2:2, 1:3, or 0:4 ratio of wild-type to SSIMS. Control matings were also performed with twenty wild-type females and three, two, or one wild-type males to account for the decrease in offspring that is due to limited wild-type male availability. Total fecundity of the females increased with each additional male from 1 to 3 males, and no significant increase with 4 wild type males in the vial (Figure 3c). Furthermore, total fecundity is reduced when SSIMS males are added to the vials, which cannot be accounted for by reducing the number of wild type males (Figure 3c).

**Fig. 3.**
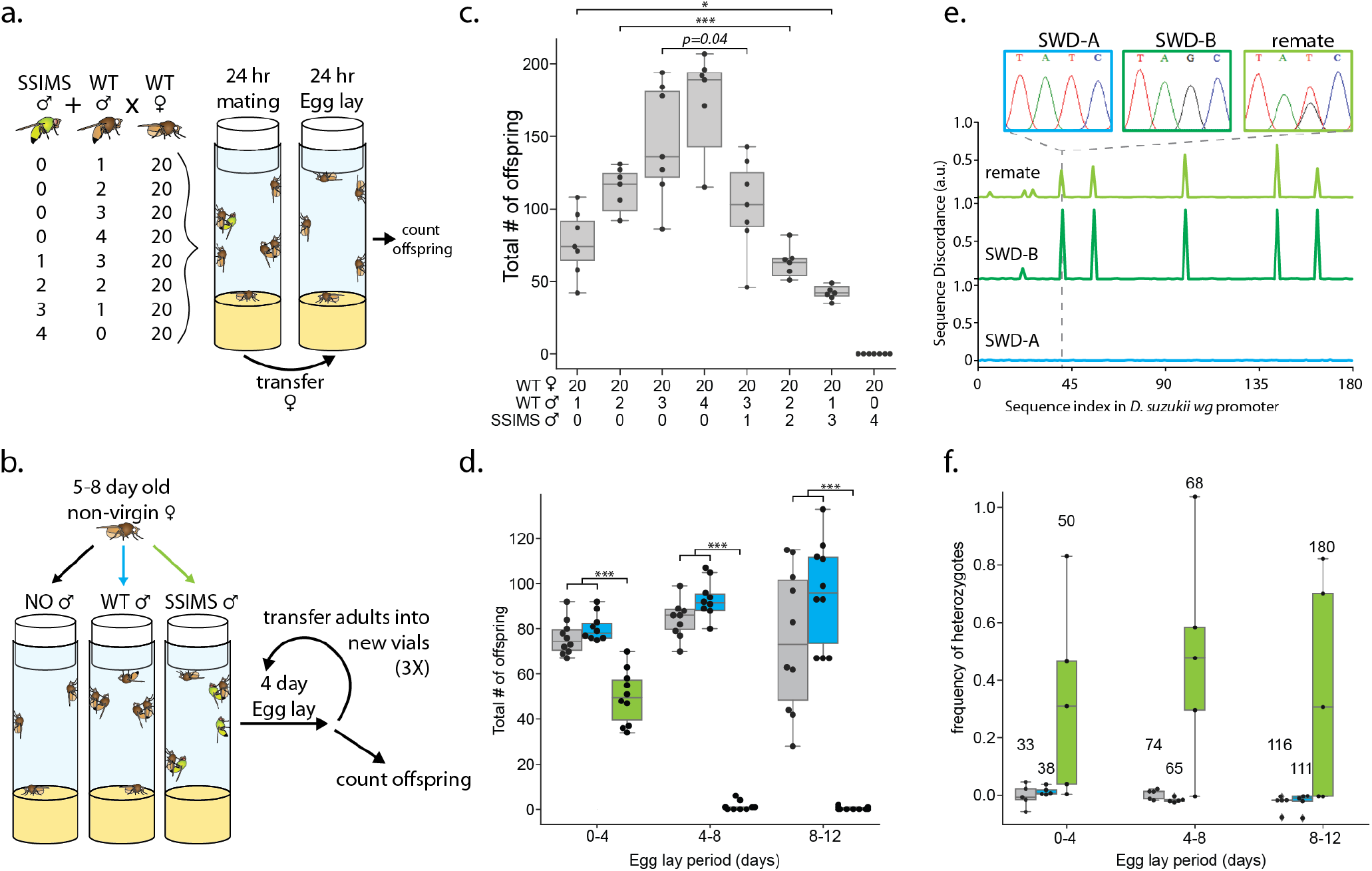
Male mating competition with wild-type *D. melanogaster*. (a) Experimental design schema and results (c) of mixed-male mating assay. Box-whisker plots show the median (center line), 25th and 75th percentile boundaries (box), and min/max (whiskers). Statistical tests of significance are performed between experiments with identical numbers of wild-type males. n=7 per group. (b) Experimental design schema and results (d) of sperm competition assay. Box-whisker plots colored to show the addition of no males (grey), wild-type males (blue), or SSIMS males (green) to the non-virgin wild-type females. Statistical tests of significance are performed for each egg-lay period comparing +SSIMS male vs both no-male and WT-male groups. n=10 per group. (e) Representative data highlighting the calculation of sequence discordance when compared to the *wg* promoter sequence of inbred strain SWD-A (blue). Inset Sanger sequence chromatographs highlight the T41G SNP in inbred strain SWD-B (dark green). The remating trace (light green) is representative of many samples from (f). (f) Heterozygote frequency when non-virgin SWD-A flies were co-housed with no males (grey), SWD-A males (blue), or SWD-B males (light green). Numbers above box-whisker plots show total number of offspring collected from each experimental group. Statistical tests of significance are performed for each egg-lay period comparing +SWD-B male vs both no-male and +SWD-A male groups. n=5 per group. Statistical significance: * = p<0.01, ** = p<0.001, *** = p<0.0001.

In *Drosophila* spp., females can store the sperm from multiple males in their spermathecae which allows for sperm to compete to fertilize the eggs (24). We tested the hypothesis that SSIMS males can reproduce with previously mated wild-type females. We placed three 5-6 day old non-virgin wild-type females (whose previous mating was with wildtype males) either with no males, three wild-type males, or three SSIMS males and measured total offspring of each group in four-day windows for a total of twelve days (Figure 3b,d). Non-virgin wild-type females with either no males or with wild-type males for re-mating, produced an average of 70-90 offspring that survived to adulthood from each vial (Figure 3d). However, when SSIMS males were available for re-mating, the fecundity of the females dropped by 36% within the first egg lay period, and by 98-99% for the next two egg laying periods (Figure 2d). In the presence of SSIMS males we observed post-zygotic lethality of eggs and larvae during all three egg laying periods, which suggests that SSIMS males mated with the females. This demonstrates that *D. melanogaster* re-mates in laboratory settings, and that the sperm from SSIMS males is competitive for fertilizing female eggs even in the presence of sperm from wild-type males in the spermatheca.

We tested whether *Drosophila suzukii* (Spotted Wing Drosophila), an agricultural pest, was similarly polyandrous in a laboratory mating assay. The experiment was similar to the sperm competition assay described above, but males from two different inbred populations were used. Strain ’SWD-A’ females were placed in vials with either no males, ’SWD-A’ males, or ’SWD-B’ males. The SWD-B population has unique fixed single nucleotide polymorphisms (SNPs) that allow us to track paternal identity of the offspring easily using Sanger sequence analysis (Figure 3e). From the first egglay period, 3 replicate had a SNP signature that suggests remating. From the second and third egg-lay period, four out of five and three out of five replicates respectively had remated (Figure 3f). These results suggests a possible important role of polyandry and sperm competition in genetic biocontrol campaigns targeting this organism.

### Population suppression cage trials

We assessed whether SSIMS could be applied to suppress wild-type populations by conducting a cage trial with multi-generational populations of wild-type *D. melanogaster* with and without additions of SSIMS animals (Figure 4a and Supplementary Note 2). In cages, adults were sustained by feeding 5% yeast extract and 5% sucrose solution via a cotton wick. To allow the population to expand, a 250 ml bottle (reproduction bottle) with 30 ml standard cornmeal food was introduced to each cage once a week for 24 hours. After 24 hours, adults were removed from the bottle and released back into the cage and the bottles were capped to prevent additional egg-lay, larvae overcrowding, and adults getting stuck in the semi-solid medium. Reproduction bottles were uncapped 10 days after egg-lay and left open for 3 days to allow for the next generation to eclose and integrate with the multi-generational caged population. To assess population size and genotype, we placed an active-yeast/sucrose trap inside each cage once per week for 24 hours. We also measured the fecundity of ten randomly captured wild-type females from each cage once per week.

**Fig. 4.**
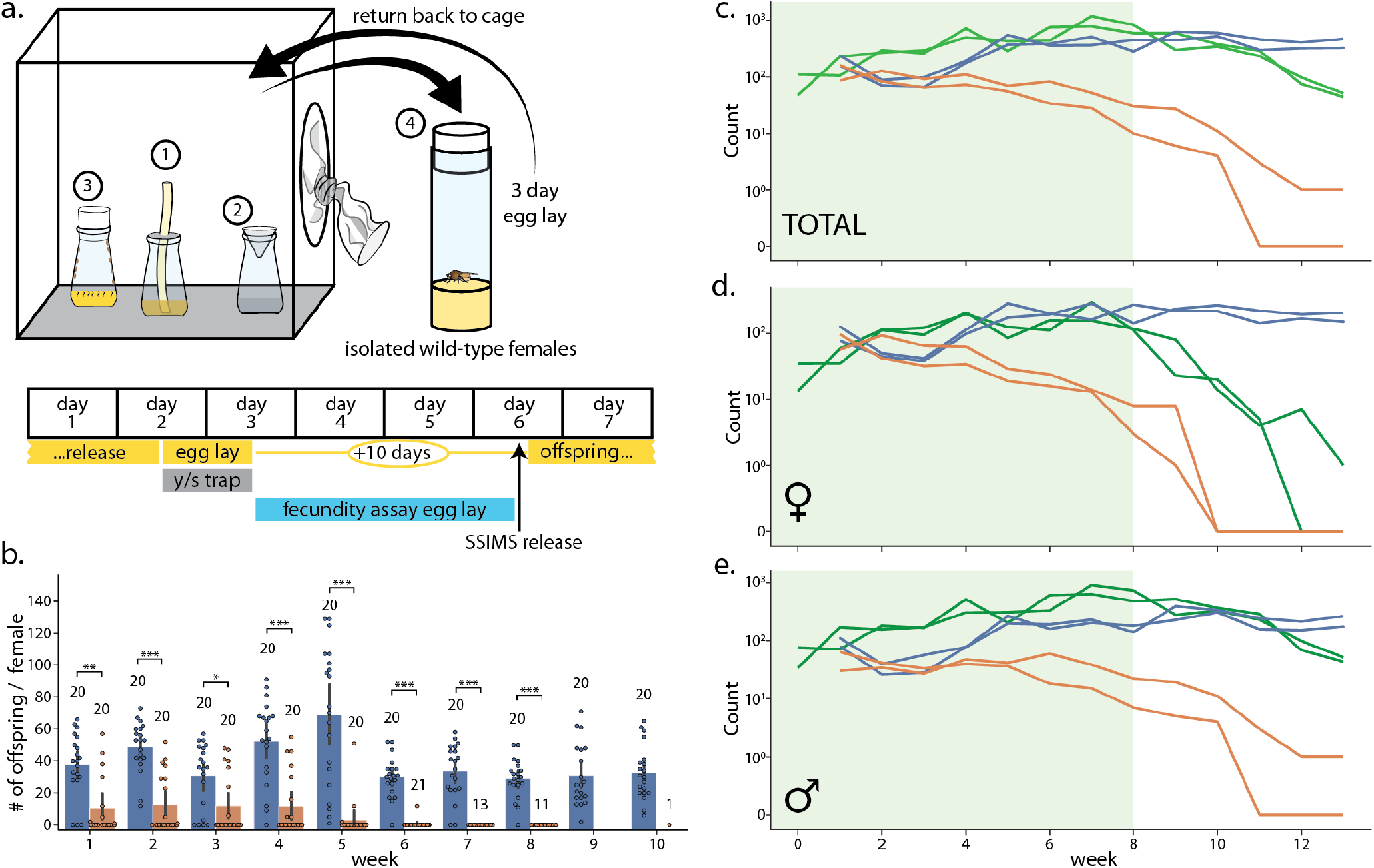
Laboratory cage trial of population suppression of wild-type D. *melanogaster*. (a) Cage trial design showing Bioquip cage with adult feeding aparatus (1), yeast/sugar trap (2), reproduction bottle (3), fecundity assay vials (4), and weekly schematic of the trial, with tasks done each day highlighted below. 2 control and 2 SSIMS treated cages were initiated, see methods for details. (b) Results of the fecundity assay from each week. Each colored dot represents total number of offspring from a single vial with one wild-type female from control cages (blue) and SSIMS treated cages (orange). Sample size of each group per week is illustrated above groups. Statistical tests of significance are performed for each week comparing control and +SSIMS treated cages, n=11-20 per group. (c-e) Total (c), female only (d), and male only (e) counts from the yeast/sugar trap. Light green shading indicates weekly additional SSIMS releases until week 8. Wild-type population in the control cages (blue lines) rise gradually and reach plateau by week 6. Wild-type population in the SSIMS treated cages (orange line) decreases gradually until no wild-type females are undetectable by week 11. SSIMS population in the treatment cages (green line) rise during weekly release period, but fall rapidly after week 8 when releases were ceased. Statistical significance: * = p<0.01, ** = p<0.001, *** = p<0.0001.

At the start of the experiment and for each of the first eight weeks, we released 600 SSIMS flies to the two experimental cages (approximately 45:55 female:male ratio) each week until week 8. These SSIMS flies had been reared on 100 *μ*g/ml Tet, so we expected the female lethality phenotype from conspecific matings to be delayed by one generation (Figure 2d-f). On week 1, all four cages were seeded with 300 wild-type virgin females and 300 wild-type males. Weekly trapping, egg-laying, and fecundity assays were performed until week 13.

The SSIMS-treatment cages saw an immediate reduction of fecundity in wild-type females. Female fecundity did not follow a normal distribution, but rather was multimodal with a mode at 0 and another mode 50, likely representing females that had mated with SSIMS or wild-type males, respectively (Figure 4b). Several females gave low numbers of offspring, possibly due to re-mating with wild-type and SSIMS males. By week seven, none of the captured wild-type females were fecund. The number of female fecundity assays that we could perform each week dropped as wild-type females could no longer be isolated from treatment cages.

Figures 4c-e show the results of the weekly trap counts. In the control cages (blue lines), counts of wild-type flies decreased initially up until week 3, at which point the weekly reproduction allowed the caged population to expand and reach a steady-state of approximately 500 flies per week (Figure 4c). While cages were too large to allow for counting of all of the individuals, we estimate by visual inspection that the total cage population was approximately three times larger than the number of trapped/counted flies.

In the SSIMS treatment cages, the counts of wild-type flies never rebounded, but dropped continuously throughout the experiment. While wild-type flies remained in each treatment cage by the time the SSIMS releases stopped in week 8, the high numbers of SSIMS males in the cages at that point prevented the wild-type population from recovering. By week ten there were no wild-type females identified in the weekly traps.

The number of SSIMS flies, both males and females, continued to increase in the trap counts during weeks one to eight. This was expected as weekly releases of SSIMS adults occurred for the first eight weeks. After SSIMS releases ended, counts of male and female SSIMS flies decreased, with female SSIMS flies decreasing rapidly to the point of disappearing as early as week 12.

## Discussion

Several new techniques for genetic biocontrol have been shown effective in proof-of-concept experiments (13, 15, 19, 25–30). Combination approaches that bring together two or more distinct techniques can exhibit synergistic behaviors that are attractive for genetic biocontrol applications. Here we combine conditional female lethality with EGI to create the SSIMS strain.

Recent advances in genetic engineering and synthetic biology have enabled the development of more complex engineered systems. The final genotype of the SSIMS flies generated in this study contains 23 synthetic genetic elements (operators, promoters, CDSs, terminators, etc.) spread across four chromosomal loci in the haploid genome. In total, 42,654 bases of synthetic DNA are integrated to the genome of engineered flies. This makes SSIMS one of the most complex engineered systems in insects.

Stacking more genetic components into a single organism can lead to fitness defects due to problems of resource allocation or cross-talk (31, 32). We observe a sex-ratio bias in favor of males when the SSIMS flies are reared with Tet, suggesting that the two X-linked female lethal constructs confer a slight fitness penalty in females even in the repressed state. This same bias was seen previously in the absence of the EGI constructs (22). While slight sex-ratio biases like this might slow the expansion of a colony during mass rearing efforts, this is likely to be offset by the cost savings attained by automatic sex-separation or by the field amplification following bi-sex release. Based on previous observations with dosage effect of the female lethal construct, we predict that fine-tuning the expression strength of the tTA-regulated promoter will provide a tuning variable to control the dynamic range of the on/off state change. Here we favored complete penetrance at the expense of genetic burden, but this should be examined more carefully in species appropriate for field applications.

The competitiveness of biocontrol males with wild-type males is a critical factor in the efficacy and economy of SIT-like approaches. We show that the SSIMS males are competitive with wild-type males in laboratory mating experiments (Figure 3a), and that this extends to the level of sperm competition. This latter point is important for polyandrous insects. If a wild female mates with both a wild male and a SSIMS male, she will have a decrease in fecundity due to this sperm competition effect. This would not occur for SIT-like approaches that prevent sperm development in males, although when females are allowed to mate with sperm-deficient males for sufficient time periods, they do appear to become satiated (33). The laboratory competition assays performed here are promising but do not necessarily predict competition in the field. Suboptimal rearing conditions (1), microbiome differences between wild and reared flies (34), or behavioral differences like assortative mating (35) could impact male mating competition. These factors should all be considered during the translation of an approach like SSIMS to field settings.

In its simplest application, SSIMS could be used to facilitate the sorting of male from female biocontrol agents prior to environmental release. In a male-only release, no biocontrol agents would persist beyond the released generation. Such an approach would be highly self-limiting, but would require larger release numbers to achieve suppression. Alternatively, males and females could be released together to provide limited field amplification. In a bi-sex release, if either sex of SSIMS insect mates with wild-type, no offspring will survive. If two SSIMS insects mate with each other, more biocontrol agents will be produced to potentiate the population suppression.

Interestingly, we can control the persistence of the SSIMS genotype to a degree by tuning the amount of Tet present in the rearing environment (Figure 2c-e). When SSIMS flies are reared in 10 *μ*g/ml Tet, female lethality occurs in the next generation (when they are reared with no Tet in the food). When SSIMS flies are reared in 100 *μ*g/ml Tet, female lethality is delayed a generation, but is still strongly penetrant. This ’tunable persistence’ could be leveraged to achieve greater population suppression with lower release numbers or frequency.

Another benefit to the SSIMS approach is redundancy. Both EGI and conditional female lethality are biocontrol techniques in their own respect. If one component were to mutate to become inactive in a released SSIMS strain, the other component would still prevent long-term persistence in most scenarios. However, this redundancy is partially reduced by the fact that the components are not linked on the chromosome. While this is a proof of concept demonstration in a model insect species, the genetic design is likely to be portable to other species for applications in pest control. The required genetic components have been shown to work in diverse organisms, and the EGI target gene for lethal ectopic expression is conserved in different insect species. The SSIMS line is true-breeding in the presence of Tet, which would facilitate mass rearing. Because the addition of Tet to rearing media would increase the cost of SSIMS applications, this will need to be factored into future techno-economic analyses, which are beyond the scope of this work.

## Conclusions

In conclusion, we present a proof of concept of SSIMS in a model insect, *D. melanogaster*. SSIMS represents a new genetic biocontrol approach that provides a unique set of strengths and weaknesses compared to other demonstrated or proposed strategies (19). In species where release of genetically engineered females is acceptable, SSIMS affords a tunable field-amplification, while still displaying low persistence. The male biocontrol agents are competitive with wild-type males for copulation and their sperm is competitive within the spermatheca. This makes SSIMS an especially attractive self-limiting genetic control approach for polyandrous pest organisms.

## Methods

### Fly stocks and rearing conditions

*D. melanogaster* strains were maintained at 25°C and 12 hour light/dark cycle and 60-70% relative humidity. All experiments were conducted in standard cornmeal/agar food Bloomington Formula (Genesee Scientific, 66-121), 0.05M propionic acid (Sigma, 402907), 0.1% Tegosept (Genesee Scientific, 20-258). Tetracycline was added to 70°C cool fly food (100*μ*g/ml, unless specified) before pouring into vials. *White1118* strain was used as a wild-type. All crosses were performed with 2-5 day old virgin females and males, unless specified.

### Generation of SSIMS line

SSIMS is generated by combining the sex-sorting female-lethal strain with the Engineered Genetic Incompatibility strains published previously (13, 22). We made a compound stock that contains 2 copies of the female-lethal construct on the X-chromosome, a refactored pyramus promoter on the 2nd-chromosome, and a programmable transcriptional activator targeting the wildtype *pyramus* promoter (dCas9-VPR)PTA+(pyr)gRNA expression cassette on the 3rd-chromosome. The combination of the refactored promoter and PTA+gRNA garners the EGI capacity of the SSIMS line. The cross strategy used to combine the two transgenes is illustrated in supplemental figure 1.

### Mating assays

Efficacy of female-lethality and EGI was measured by crossing 2-5 day old SSIMS male and virgin females (SSIMS and wild-type) with or without Tet (10*μ*g/ml). 5 virgin females and males were were allowed to mate in vials for 3 days, then removed to prevent overcrowding of larvae. Male and female offspring were counted 10 days post egg lay.

### Male competition assays

To assess SSIMS male competitiveness various combinations of 0-4 SSIMS and wild-type males were placed together with 20 wild-type virgin females, and allowed to mate for 24 hours. After the mating period, males were discarded and females were transferred into a new vial for 24 hour egg lay period. Total offspring were counted 11 days post egg lay.

### Sperm competition assay

To assess the ability of SSIMS sperm to compete with wild-type sperm *in utero*, 3 (5-8 day old) non-virgin wild-type females, which were previously mated with wild-type males, were placed in a vial either alone, with 3 wild-type male, or 3 SSIMS male. Females (with or without males) were allowed to lay eggs in 4 day periods then flipped into new vials for a total of 12 days. Total offspring were counted 11 days post egg lay for each period. To investigate polyandry and sperm competition in *D. suzukii*, two inbred SWD lines, with unique fixed SNPs in the 5’ promoter region of the *wg* gene, were crossed. Mating pairs of 3 non-virgin ’SWD-A’ females with 0 or 2 males of ’SWD-A’ or ’SWD-B’ were placed in vials. Genomic DNA of adult offspring were isolated as previously described (22). PCR of the *wg* promoter region was done using primers wgF (5’-AGATTGCGCAAATAATCCGGC-3’) and wgR (5’-ATTCGAGCGGAGGAGTGAAG-3’), and Q5 polymerase following manufacturer recommendations(New England Biolabs). PCR products were Sanger sequenced by ACGT (Wheeling, IL), and the results were analyzed by Synthego’s ICE software (ice.synthego.com).

### Cage trial

Cage trial experiments were designed to simulate multi-generational and generational-overlapping populations dynamics of wild-type *Drosophila*. On week 1, the cage trial was initiated with 600 (50% female) wild-type virgin males and females into each cage. 600 SSIMS adults (40-50% female) were released weekly in the +SSIMS treatment cage, and once before wild-type release. Adult populations were housed in 0.027 m^3^ cages (Bioquip, 1452) and were fed by supplying 50ml of 5% yeast-extract, 5% sucrose (YES), 0.1% Tegosept solution via a cotton wick in a 250ml standard fly bottle and wrapped at the rim to prevent flies from getting stuck in the bottle. YES solution was replaced weekly. To allow the population to grow, 250ml bottle (reproduction bottle) with 30ml standard CMF were introduced once a week for a 24 hours egg lay period, after which the adults were removed. Reproduction bottle were uncapped 10 days after egg-lay and left open for 3 days to allow for the next generation to eclose and mix/mate with the preexisting population in the cage. To assess population size and genotype, an adult fly trap with yeast/sugar (YS) solution was placed in the cage once a week for 24 hours. The fly trap was baited with 30 ml of YS solution consisting of 0.5% active-yeast, 2.5% sucrose, 0.5% Micro90 (Cole Parmer). To measure fecundity of wild-type females, once a week, random adults from each cage were trapped in CMF bottle and wild-type females were isolated individually into 10 separate vials, and the remaining adults were released back in the cage. Wild-type females were allowed to lay eggs in vials for 3 days then released back in the cage. Total offsprings from each vial was quantified 11 days post egg-lay.

### Data analysis, statistics, and visualization

All experiments were performed at least twice except the cage trial. Each batch of experiments contained at least 5 replicates, see caption for each figure. Chi-squared test was perfomed to test difference between observed and expected sex ratio for figure 2b and student’s t-test for figure 2c. For figure 3, a one-way ANOVA with Tukey post hoc test for multiple comparisons. For figure 4b, Mann-Whitney non parametric test was applied. Data was analyzed in Python using pandas, scipy, and statsmodels libraries. Statistical significance: * = p<0.01, ** = p<0.001, *** = p<0.0001. Graphs were generated using the Seaborn and Matplotlib libraries. Graphs were modified in Illustrator to fit publisher requirements.

## ACKNOWLEDGEMENTS

We would like to thank Dr. Max Scott for helpful discussions about conditional female lethality and for providing us with genetic reagents. We would like to thank Dr. Michael O’Connor for balancer strains that were used for the cross strategy. MJS, AU, SD, and MM were supported in part by the Defense Advanced Research Projects Agency (grant number D17AP00028). The views, opinions, and/or findings contained in this article are those of the authors and should not be interpreted as representing the official views or policies, either expressed or implied, of the Defense Advanced Research Projects Agency or the Department of Defense. AU and NF were supported by a grant from the Minnesota ENRTF to the Minnesota Invasive Terrestrial Plants and Pests Center.

## Supplementary Note 1: Mating strategy to produce SSIMS line

**Supplemental Figure S1.**
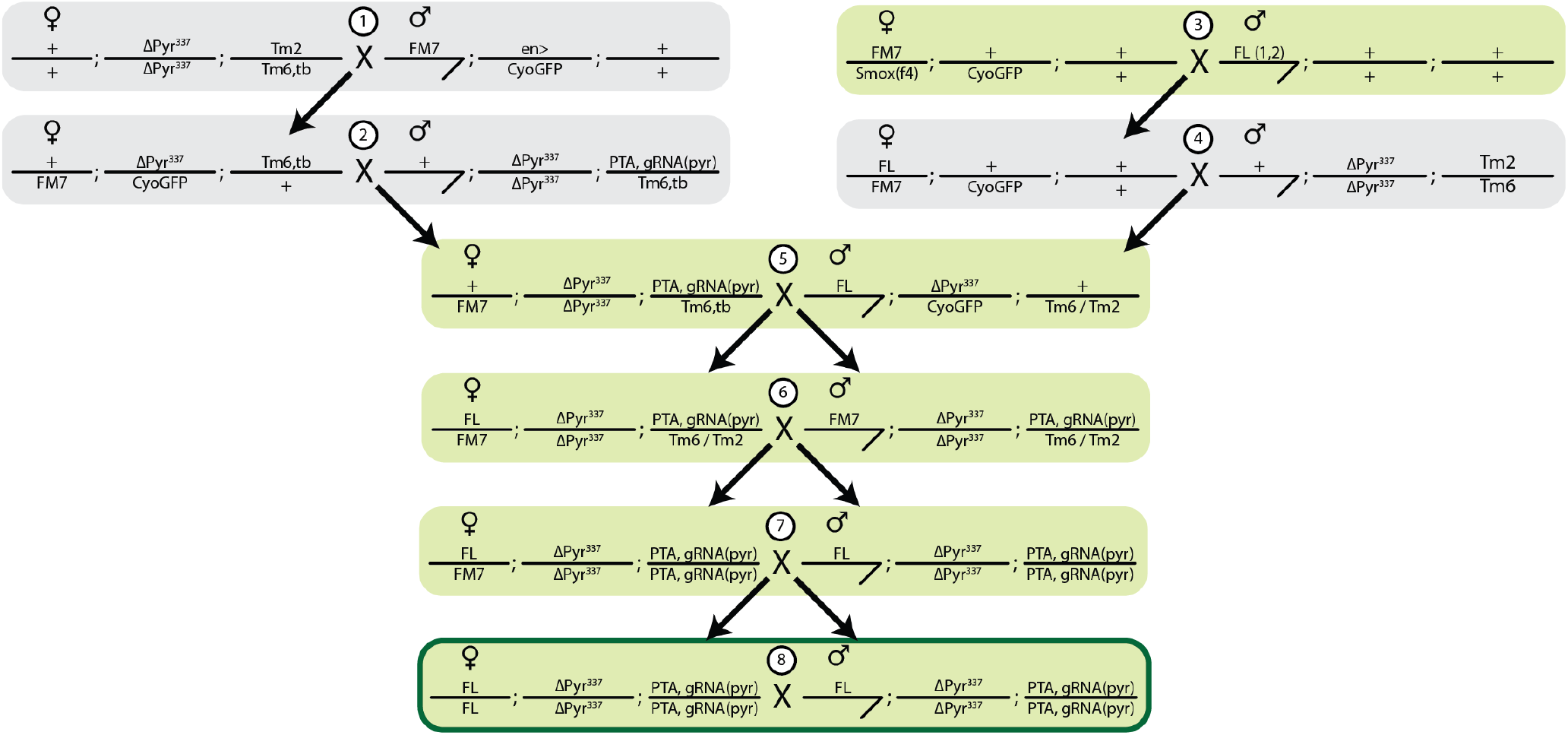
Cross strategy to generate SSIMS line. Female lethal (FL) and EGI (pyr337; PTA, gRNA-pyr) were crossed to balancer chromosomes, and subsequent males and females were isolated indicated by the black arrow. FL chromosomes contains 2 insertions on the X chromosomes, homozygous females contain 4 copies of the transgene. PTA+gRNA-pyr is a single insertion site containing foxo driven dead-Cas9::VP64 and 2 guide RNAs driven by U6:3 and U6:1 promoters. Green shading indicates crosses done in the presence of 100*μ*g/ml Tet. FL was tracked by GFP, PTA+gRNA was tracked by red eyes (mini white), and the pyr337 promoter mutation was tracked using balancers.

## Supplementary Note 2: Extended timeline of cage trial design

**Supplemental Figure S2.**
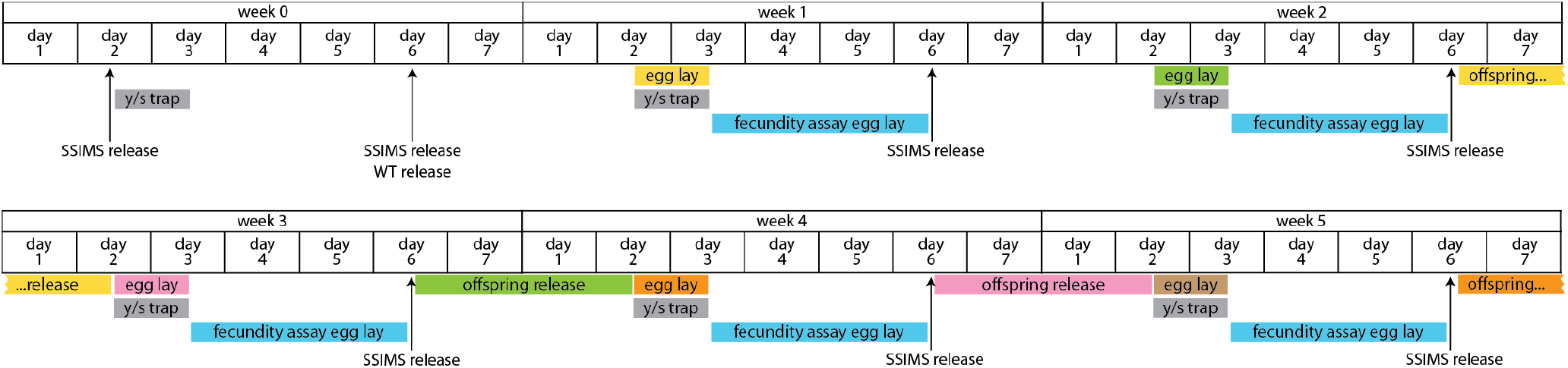
Extended timeline of cage trial design. In this multi-week view of the cage trial data, the timing between egg lay and offspring release is shown by coloring corresponding events. During the ten days between egg lay and offspring release, the bottles were kept in cages, but were capped with a foam plug to prevent caged flies from entering and becoming stuck in the cornmeal food.

## Supplementary Note 3: Sample pictures of female fecundity assay and reproduction bottles during large cage trials

**Supplemental Figure S3.**
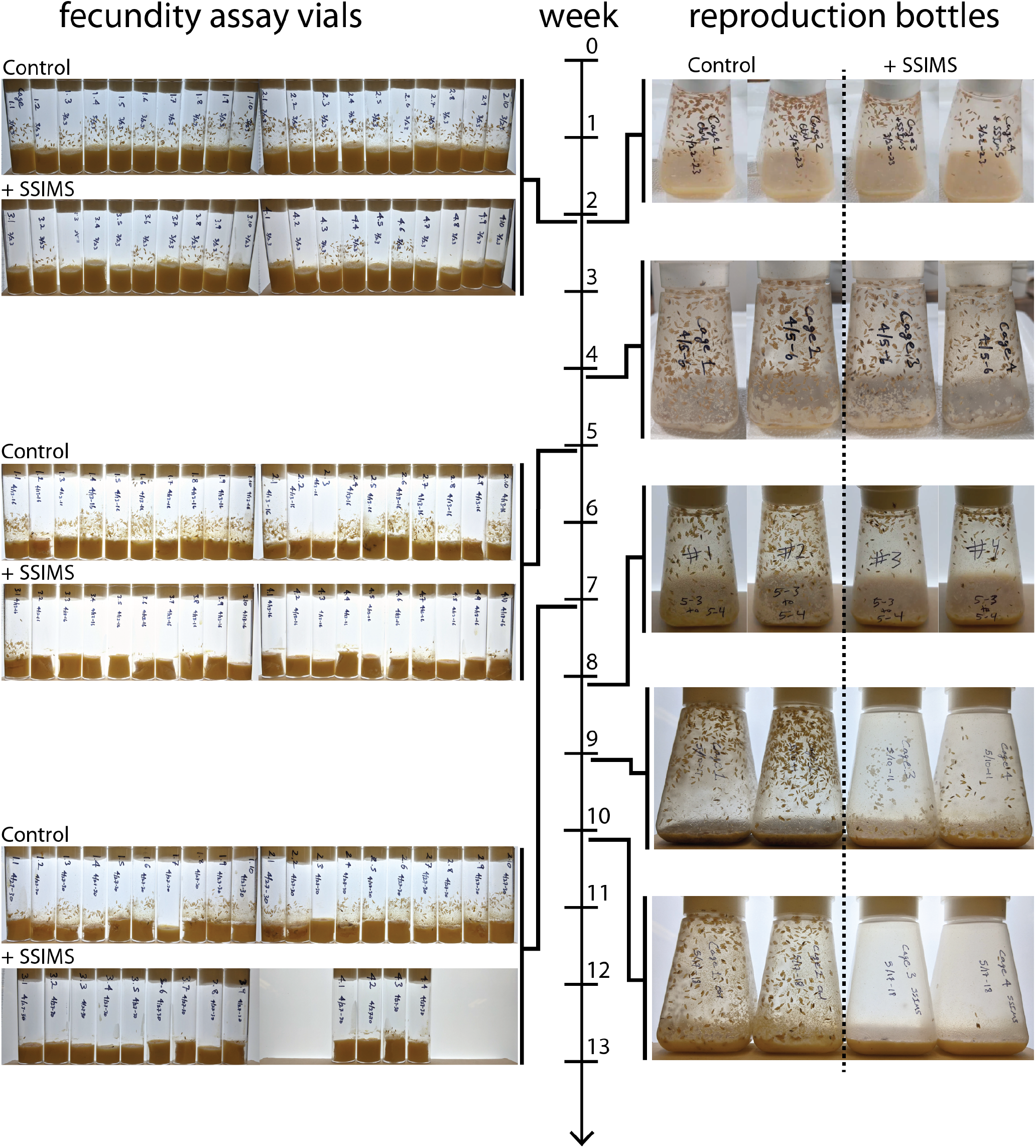
Sample pictures from cage trial. Timeline of cage trial shown in middle. Jagged lines indicate the week when images were taken. Fecundity assay vials from control cages and SSIMS treated cages to the left of the timeline. Images of the reproduction bottles from the control and SSIMS treated cages to the right of the timeline.

